# Sniffing Shapes Dopamine Signals for Reward Prediction

**DOI:** 10.64898/2026.04.27.721190

**Authors:** Max F. Scheller, Catarina Luís, Wolfgang Kelsch

## Abstract

Adaptive behaviors depend on predicting outcomes from sensory evidence. Dopamine neurons in the ventral tegmental area (VTA) broadcast reward-prediction signals that guide learning. Yet the principles functionally coordinating information flow from input regions to VTA are incompletely understood. In the olfactory system, the sniff cycle structures sampling and odor encoding. We therefore asked whether this rhythm also entrains the ventral striatum to VTA communication and if so, how this shapes the implementation of predictive coding in dopamine neurons. We recorded identified dopamine neurons throughout olfactory conditioning and found that their firing shifted systematically to the post-inspiratory phase of the sniff cycle with learning. This temporal realignment predicted a neuron’s engagement in value encoding along the optimism-pessimism-spectrum of distributional reinforcement learning. This is associated with an enhanced phase-gated communication channel from the striatal olfactory tubercle to dopamine neurons, the strength of which predicts task performance. Thus, the sniffing rhythm provides a scaffold for information flow, revealing a phase-gating mechanism for the integration of outcome predicting sensory evidence to dopamine neurons during reinforcement learning.

**Significance Statement:** Predicting the future from sensory cues is central to adaptive behaviors. We show that sniffing temporally organizes the flow of information between the olfactory tubercle of ventral striatum and midbrain dopamine neurons during odor-guided learning in mice. Phase-specific coupling in the respiratory cycle determines when sensory information reaches dopamine neurons and how predictive signals are encoded. These findings link active-sensing rhythms to predictive reinforcement signals. This temporal scaffold yields a gradient of “optimistic” to “pessimistic” predictions consistent with distributional reinforcement-learning theories. These insights contribute to a better understanding of the large-scale computations across multiple brain regions to predict future outcomes.

## Introduction

The mesolimbic dopamine system is fundamental for reinforcement learning. Phasic activity of dopamine neurons (DANs) in the ventral tegmental area (VTA) signals reward prediction errors, i.e. the difference between expected and actual outcomes (1). Upon learning, sensory cues like an odor that reliably predicts a reward, will elicit a predictive response in DANs (2, 3).

To encode these predictions, the VTA DANs integrate information from numerous brain regions (4). Prominently, these input regions involve the ventral striatum, that is composed of two subregions, namely the nucleus accumbens and the olfactory tubercle (Tu) (5, 6). For olfactory guided behaviors, the Tu is uniquely positioned at the interface of sensory and reward processing regions, receiving direct input from the olfactory bulb and cortices (7). At the same time, the Tu receives some of the densest levels of innervation from VTA DANs (7–9). In this bidirectional VTA-Tu loop, dopaminergic signaling reinforces odor-evoked responses in the Tu thereby generating sensory-driven expected outcome coding (10–13). In turn, the Tu can inform the predictive signaling of VTA dopamine neurons (13).

The classical account of dopamine firing signaling the expected value of a stimulus has been extended in recent years. Research in deep reinforcement learning revealed a class of algorithms that result in higher performance for agents using the return distribution as a learning target instead of the scalar expected return (14, 15). This distributional representation allows, for example, to distinguish between stimuli with different “risk”, but the same expected return, presumably a distinction with relevance also for biological agents. Indeed, it was found that DANs exhibit heterogeneous scaling of prediction errors consistent with learning different expectiles of the return distribution(16). A first foray into how this representation may arise in DANs was made in a recent study that proposed a segregation by dopamine receptor dependent learning in striatal projections neurons (SPNs) (17).

Relevant to the biological implementation of reinforcement learning, it has remained an open question how the VTA-Tu loop is coordinated to generate predictive signals during the course of learning. It has been shown that phasic dopamine in the Tu reinforces conditioned stimulus (CS) responses (11–13), yet the principles organizing the observed information flow from ventral striatum to VTA remain incompletely understood. Specifically, how is the long-range communication from Tu to VTA coordinated to generate odor-triggered predictive responses in DANs? Thus, we aimed to identify the mechanism that orchestrates the activity of neuronal ensembles across these interconnected regions as animals learn.

Neural oscillations can coordinate such long-range communication. The communication-through-coherence (CTC) hypothesis posits that synchronized oscillations create temporally aligned windows of excitability, facilitating effective communication between neuron groups (18, 19). Both striatal (20, 21) and VTA (22–24) neurons phase-lock to local field potential oscillations. Here, we propose that sniffing, i.e. the inhalation-exhalation cycle, may provide a key organizing rhythm also to the mesolimbic system and in particular to DANs. Sniffing not only structures olfactory primary sensing (25–27) and secondary processing in the olfactory network (28, 29), it also entrains widespread neural oscillations (30), acting as a global temporal framework for neural processing that has been shown to influence perception and cognition (31–34), with evidence for behavioral state-dependent differential sniff-phase coupling of single units even in the medial prefrontal cortex (35). In the context of olfactory learning, dopamine and sniffing are linked, with dopamine transients in the ventral striatum modulating sniff frequency (36). However, the reverse influence, i.e. the influence of the sniffing rhythm on DANs, has not yet been studied.

We hypothesized that the sniff cycle may provide a temporal scaffold that organizes DAN responses during odor-reward association learning. We asked whether VTA DANs firing is aligned to the sniff cycle as animals learn to predict reward, and if alignment is related to the encoding of predicted value and its features in a distributional reinforcement learning framework. Furthermore, we investigated whether sniff-phase coupling facilitates directed communication from Tu to VTA, in line with the CTC framework, and how this information channel is related to the animals’ behavior.

## Results

### Respiratory Rhythm serves as a Carrier for DAN activity

To investigate the potential interplay between sniffing, learning, and dopamine signaling, we recorded simultaneously from optogenetically identified dopamine neurons (iDANs) in the VTA and putative striatal projection neurons (SPNs) in the Tu of head-fixed mice in a custom-built olfactometer (Fig. 1A, Fig. S1, Methods). Mice were trained on a Go/No-Go task where they learned to associate three different odors (conditioned stimuli, CS) with varying reward probabilities (100%, 50%, or 0% reward, referred to as CS_100%_, CS_50%_ and CS_0%_; Fig. 1B, C). This discrimination learning paradigm was preceded by a phase of behavioral shaping, where only one odor with 100% reward probability was presented. In both learning phases, behavioral performance was defined as the fraction of trials where animals reported their anticipation by licking before reward delivery in Go trials or withheld anticipatory licking in No-Go trials. All animals achieved criterion performance (80% correct trials) for at least 3 sessions in the discrimination paradigm, although learning speed and performance reliability varied between individuals (Fig. 1C, D). As expected, iDANs (Fig. 1F, Methods) encoded the learned value of the odors after training and showed graded responses at CS that corresponded to the reward probability associated with the respective odor (Fig. 1G).

**Figure 1.**
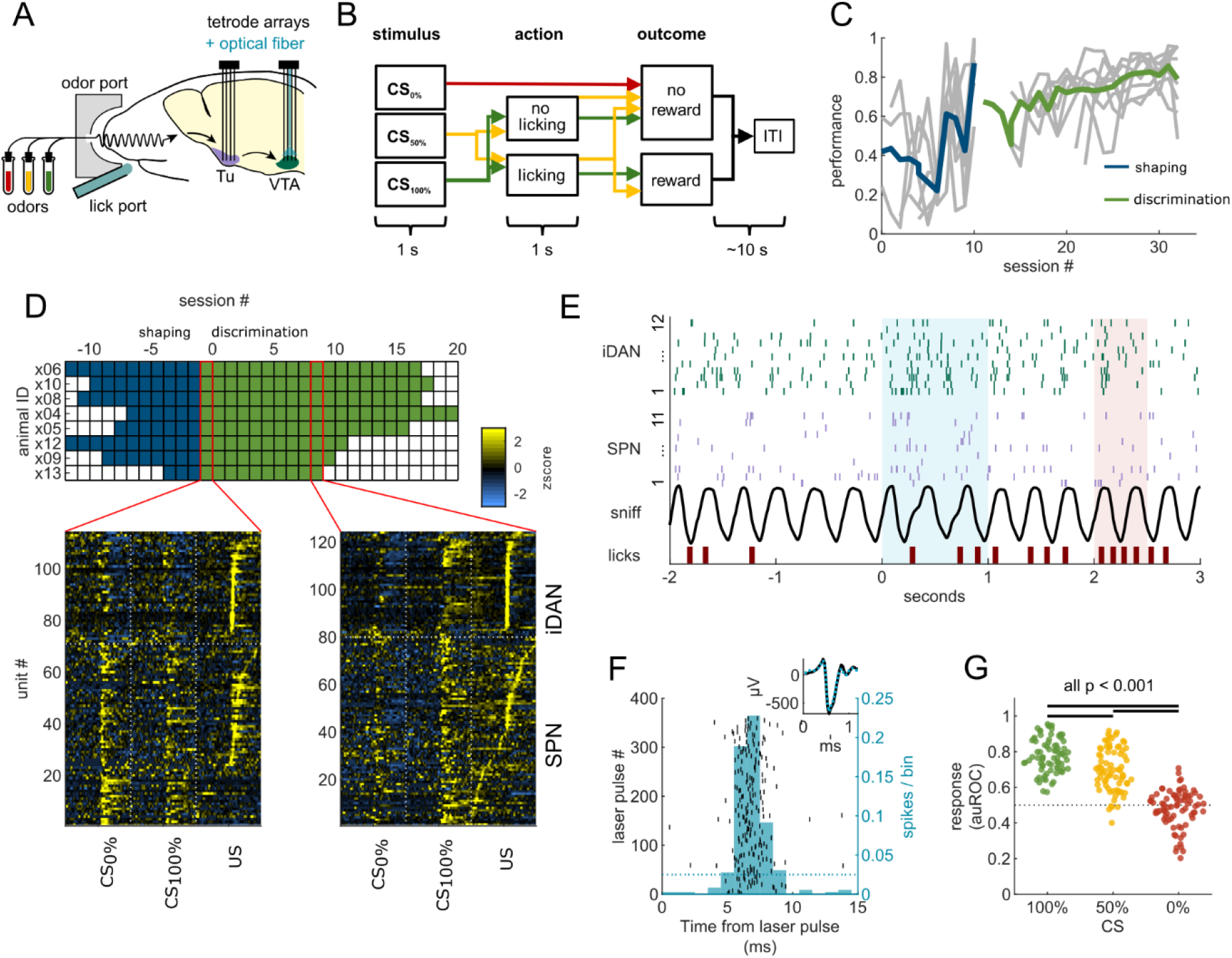
Training and recording mice in a Go/No-Go reinforcement learning task. **(A)** Schematic of the experimental setup. Mice were implanted with tetrode arrays in the VTA and Tu. Sniffing and licking were recorded in a head-fixed configuration while water rewards and different odors were presented. **(B)** Schematic of the behavioral paradigm. Mice were trained on a Go/No-Go task with three odors predicting different reward probabilities (100%, 50%, 0%). Discriminatory performance was assessed by anticipatory licking recorded after odor offset and before reward delivery. **(C)**Behavioral performance across eight mice for the initial shaping phase (single odor) and subsequent discrimination learning (three odors). Performance is defined as the fraction of trials with correct Go or No-Go response. Individual mice are shown in gray; population averages are shown in blue (shaping) and green (discrimination). Sessions are aligned to the last shaping session. **(D)** Top: Overview of recording sessions across the different contingencies for all animals. Bottom: Heatmaps of normalized firing rates (z-score) for simultaneously recorded SPNs and iDANs on two representative recording days (early and late in conditioning). Activity is aligned to trial events (CS: conditioned stimulus; US: unconditioned stimulus). **(E)** Example trial showing simultaneously recorded spikes from iDANs (green ticks) and SPNs (purple ticks), alongside the respiratory sniff trace (black) and lick events (red bars). The blue shaded area corresponds to stimulus presentation (1 s), and the red shaded area to reward delivery. **(F)** Example of optogenetic identification of a DAN. The raster plot (black) and peristimulus time histogram (PSTH, blue) of spike times aligned to a laser pulse show reliable, short-latency activation. The dotted line represents the 97.5th percentile of the null distribution generated by laser time dithering (±60 ms, 1000 iterations). Inset: Overlay of the mean spontaneous action potential waveform (black) and the laser-triggered average waveform (blue dotted line), confirming isolation stability. **(G)** Response strength (area under the receiver operating characteristic curve, auROC) of all recorded iDANs from expert animals to the presentation of odors associated with 100%, 50%, or 0% reward probability. Each dot represents one neuron. Repeated measures ANOVA (*F*(2, 152) = 239.02, *P* < 10^−45^) followed by Tukey’s post-hoc tests (all *P* < 0.001).

We first asked whether the firing of iDANs was modulated by the ongoing sniff cycle. We observed that even during the inter-trial interval (ITI), the firing of most iDANs was significantly coupled to the sniff cycle (example iDAN shown in Fig. 2A). This intrinsic coupling was present from the earliest training sessions and remained robust throughout learning, with both the fraction of coupled neurons and the strength of modulation staying consistently high (Fig. 2B).

**Figure 2.**
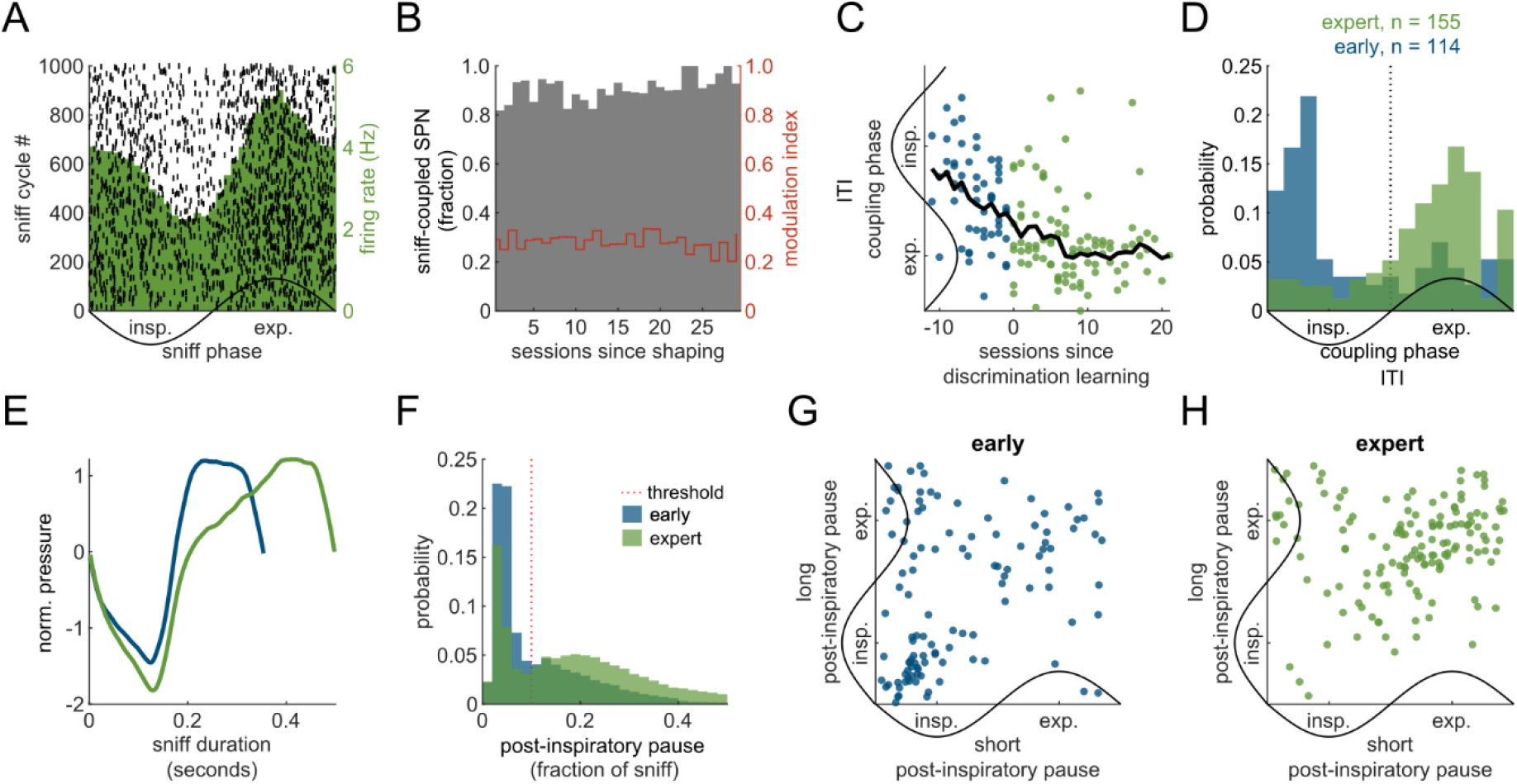
iDANs are coupled to the sniff cycle and their coupling phase shifts with learning. **(A)** Sniff-aligned peristimulus time histogram (PSTH, green) and raster plot (black) of an example iDAN during the ITI, demonstrating robust modulation of firing rate by the phase of the sniff cycle. Inspiration (insp.) and expiration (exp.) phases are indicated. **(B)** *Left y-axis:* Fraction of iDANs significantly coupled to the sniff cycle during the ITI across learning sessions (see Methods). *Right y-axis:* Mean modulation index for the coupled population (red line). **(C)** Preferred coupling phase of iDANs during the ITI for each session, plotted against days from the start of conditioning. The black line represents a moving average. Note the systematic shift from early phases (inhalation) to later phases (expiration) as training progresses. **(D)** Distribution of preferred coupling phases for all iDANs during early (blue; first 3 sessions) and expert (green; last 3 discrimination sessions) learning phases. The distribution shifts significantly towards later phases in expert animals (Watson’s *U*^2^ test, *U*^2^ = 2.19, *P* < 10^−18^). **(E)** Representative sniff traces from an expert animal, illustrating the distinction between sniffs with a short (blue) and long (green) post-inspiratory sniff pause. **(F)** Distribution of the duration of the post-inspiratory sniff pause (expressed as a fraction of total cycle duration) in early (blue) vs. expert (green) sessions. Expert animals show a significantly greater proportion sniffs with long pause (χ^2^-test; *N* early = 15,189, *N* expert = 28,706 sniffs; χ^2^ = 3540, *P* ≈ 0). **(G–H)** Circular distribution of iDAN coupling phases derived separately from sniffs with long or short post-inspiratory pauses, shown for the early training **(G)** and in expert mice **(H)**. The learning-dependent shift from inhalation to post-inspiration/expiration occurs independently of the specific sniff waveform.

While prevalence and strength of sniff-coupling in iDANs during the ITI was largely constant, we found that the temporal nature of this coupling changed with learning. In early training sessions, iDANs preferentially fired during the inhalation phase of the sniff cycle (Fig. 2D, blue). However, as animals became proficient at the task, the preferred firing phase systematically shifted to the post-inspiratory pause and early exhalation (Fig. 2C, D).

We considered whether this shift could be explained by changes in sniffing behavior itself. With training, mice exhibited a greater proportion of sniffs with a prolonged post-inspiratory pause (Fig. 2E, F). To dissociate the neural changes from the behavioral changes, we analyzed the preferred firing phase separately for sniffs with short versus long plateaus. We found that the learning-dependent shift from inspiratory to post-inspiratory/expiratory phase-coupling occurred largely independent of the trial-by-trial variations in the length of the post-inspiratory pause (Fig. 2G, H). Only a small fraction of neurons (16.1 % in early, 12.5 % in trained) showed a shift from inspiration to expiration in sniffs with a longer post-inspiratory pause. This supports that the change in neural coupling represents a genuine reorganization of iDANs to sniff coupling largely independent of the modifications in the shape of the sniff cycle.

Taken together, these findings suggest that respiration serves as a “carrier signal” on which iDAN activity is temporally organized even in resting state recording sessions in animals naïve to the stimulus-outcome associating nature of the experimental task (Fig. S2).

### Respiratory Entrainment Underlies Predictive Value Coding

Next, we investigated the functional significance of the systematic learning-dependent phase shift. The fraction of iDANs responding to task events, namely the CS and the unconditioned stimulus (US) increased as animals approached criterion performance (Fig. 3A). We hypothesized that the ITI coupling phase might determine a neuron’s readiness to participate in coding for the predictive value of the CS. We next assessed the ability of iDANs to discriminate between the different reward-predicting odors depending on their preferred ITI coupling phase. In untrained animals, iDANs did not discriminate between the three stimuli (Fig. 3B, blue). In trained animals, iDANs only significantly discriminated CS value if they preferentially coupled to the post-inspiratory ITI phase (Fig. 3B, green).

**Figure 3.**
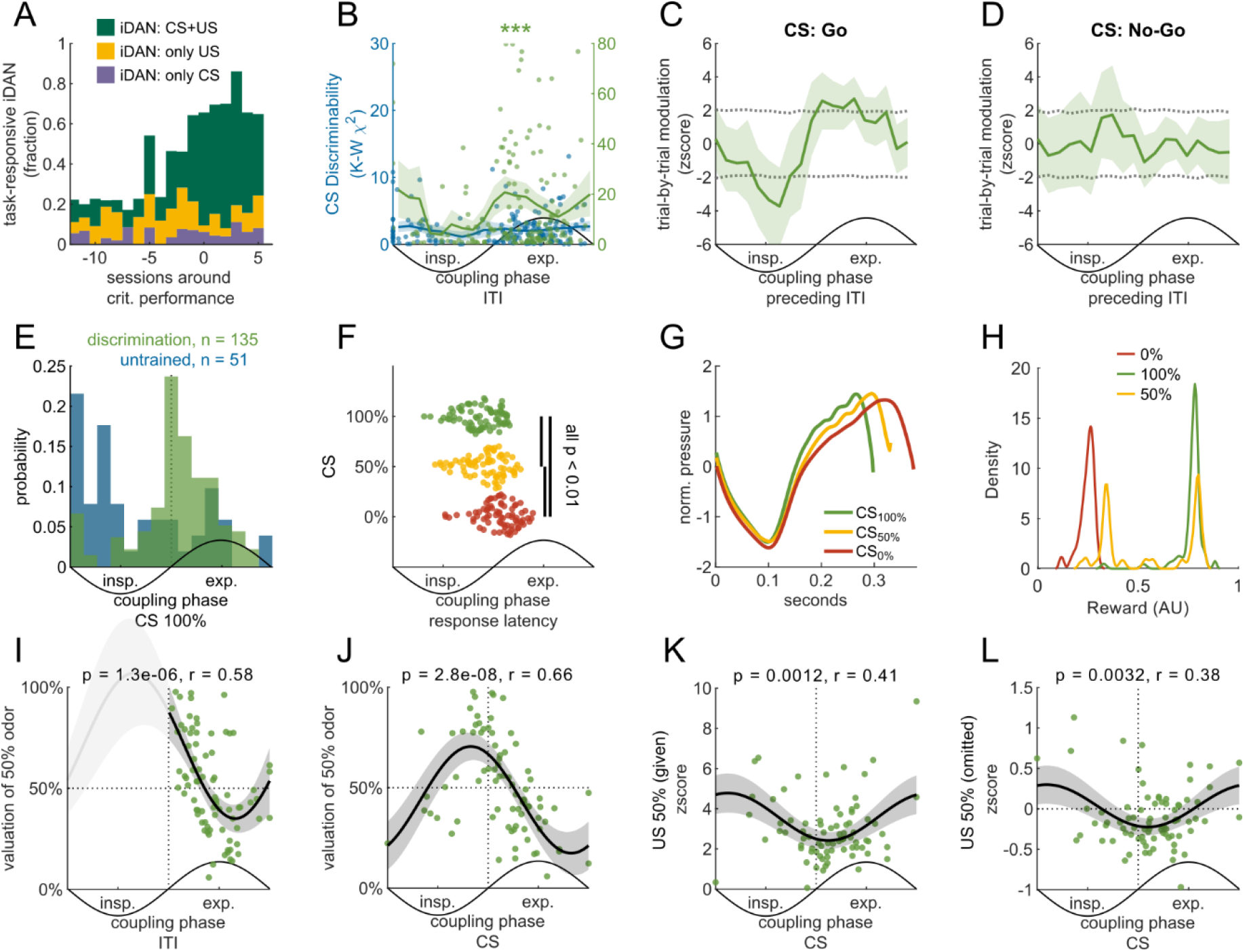
Sniff-phase coupling of iDANs predicts and organizes predictive stimulus coding. **(A)** Fraction of task-responsive iDANs across sessions, aligned to the day criterion performance was reached (day 0). Neurons are categorized as responsive to the CS only (purple), US only (yellow), or both (green) based on Wilcoxon signed-rank test vs. baseline. **(B)** CS discriminability (Kruskal-Wallis χ^2^ statistic) as a function of the neuron’s preferred ITI coupling phase. In trained animals (green, *n* = 166 iDANs), discriminability is significantly modulated by phase, peaking in the post-inspiratory/expiratory phase (*P* = 0.004, permutation test). No significant modulation is seen in untrained animals (blue, *n* = 184). **(C–D)** Trial-by-trial modulation of CS responses by the coupling phase observed in the immediately preceding ITI in trained animals. The plots show modulation of the z-scored CS response as a function of the pre-stimulus ITI phase for Go trials (*n* = 121 units, *P* = 0.003, permutation test) **(C)**, but not for No-Go trials (*n* = 155 units, *P* = 0.92) **(D)**. **(E)** Distribution of preferred coupling phases during the CS response for all iDANs in early (blue) vs. expert (green) sessions. The response phase shifts significantly to later in the sniff cycle with learning (Watson’s *U*^2^ test, *U*^2^ = 1.09, *P* < 10^−9^). **(F)** Latency of the first spike burst during the CS presentation for the three reward contingencies in expert animals. Latency is shorter for higher-value odors. Repeated measures ANOVA (*F*(2, 152) = 56.85, *P* < 10^−18^) followed by Tukey’s post-hoc tests (all *P* < 0.01). **(G)** Average aligned sniff waveforms for CS with different reward contingencies (CS_100%_, CS_50%_, CS_0%_) in expert animals. Inspiratory duration becomes shorter with higher value. **(H)** Using iDANs’ CS responses to decode the probability density over reward shows a bimodal distribution for CS_50%_ corresponding to the two possible outcomes, consistent with distributional reinforcement learning (see Methods). The distributions for CS_0%_ and CS_100%_ are plotted for reference. **(I–L)** Relationship between coupling phase and measures of value encoding in iDANs of expert animals (*n* = 80) in CS_50%_ trials. Each dot represents a neuron. **(I)** Correlation between ITI coupling phase and the normalized valuation during the CS presentation (circular-linear correlation, *r* = 0.58, *P* = 1.3 × 10^−6^). This reveals the placement of iDANs along the optimism–pessimism axis for stimulus evaluation. **(J)** Same as in **(I)**, but using the phase coupling during CS_50%_ presentation (*r* = 0.66, *P* = 2.8 × 10^−8^). Neurons switching from expiratory to inspiratory coupling extend the optimism–pessimism axis during inspiration, with earlier responding neurons acting more “pessimistic.” **(K–L)** iDANs coupled to the post-inspiratory phase also behave more “optimistically” during the outcome window. Panels show the correlation between the CS response phase and the response to a delivered **(K)** or omitted **(L)** reward for the CS_50%_ (delivered: *r* = 0.41, *P* = 0.0012; omitted: *r* = 0.38, *P* = 0.0032).

This relationship also held on a trial-by-trial basis. For individual neurons, the coupling phase during the ITI immediately preceding stimulus presentation significantly modulated the neuron’s subsequent response to the rewarded odors (Fig. 3C), but not to the unrewarded odor (Fig 3D). The modulation was also absent in untrained animals (Fig. S3A). Together, these findings suggest that the learning-induced shift towards a post-inspiratory coupling phase functionally configures iDANs to engage in predictive coding.

Furthermore, we found that the CS response itself is temporally organized by the sniff cycle. In trained animals, the peak of the value-encoding CS response occurred at a later phase within the first sniff after odor onset compared to the response in naïve animals (Fig. 3E, see also Fig. S3B, C for CS_50%_ and CS_0%_). We observed that the latency of the first spike burst after odor onset also encoded value (Fig. 3F). This latency difference can be partly explained by a shorter inspiration phase for higher value odors (Fig. 3G). In line with the parallel phase switch in both ITI and during stimulus presentation, the coupling phase of individual iDANs was highly correlated between the ITI and the stimulus presentation (Fig. S3E). In summary, the phase-switch to post-inspiration during both ITI and stimulus marks the switch of iDANs from (presumably) stimulus detection to stimulus evaluation.

We next asked whether phase coupling was related to how individual iDANs evaluate different reward probabilities. In the framework of distributional reinforcement learning, individual iDANs do not code for the average expected value, but each neuron has a position on a spectrum of ‘optimistic’ to ‘pessimistic’ evaluations. For CS_50%_, an optimistic neuron might fire as if a reward is certain, while a pessimistic one fires as if no reward were to be expected. This heterogeneity allows the population to represent the full distribution of possible outcomes, not just the mean (15, 16). To test whether the heterogeneity of iDANs’ CS responses allows to reconstruct the reward distribution associated with each odor in our data, we applied the distribution-decoding approach described by Dabney et al. (16) (see Methods for details). This yielded a bimodal distribution for CS_50%_ (Fig. 3H), corresponding to the two expectation outcomes, one being more close to the expected outcomes CS_0%_ and the other peak being closer to the expected CS_100%_ outcome, respectively; thus supporting that the heterogeneity in CS responses reflects distributional coding rather than noise.

Next, we extended our analysis to the contribution of individual iDANs. We found that a neuron’s position on the optimistic-pessimistic spectrum was systematically predicted by its preferred firing phase during the ITI (Fig. 3I): neurons coupled to the post-inspiratory phase were more optimistic, while those coupled to the later expiratory phase were gradually more pessimistic, with the average being indistinguishable from the expected 0.5 (bootstrapped CI = 0.46-0.56). When looking at the sniff coupling of the CS response itself, a similar picture emerged (Fig. 3J), although iDANs with shifted coupling to inspiratory phases were more ‘pessimistic’ the earlier their response peaked during inspiration. The principle further extended to the US time point, where a neuron’s CS response phase predicted how it coded for both delivered and omitted rewards following the uncertain CS_50%_ (Fig. 3K, L). Specifically, neurons coupled to the post-inspiratory phase showed weaker responses to delivered rewards and stronger suppression to omitted rewards, consistent with a higher initial prediction (‘optimism’).

In summary, learning reconfigures the temporal dynamics of iDANs in two key ways. First, a shift in baseline sniff-phase coupling to the post-inspiratory phase appears to prime neurons to participate in predictive coding. Second, the sniff cycle provides a temporal scaffold for representing reward uncertainty. We find that a neuron’s preferred firing phase systematically maps onto its identity within the distributional reinforcement learning framework, organizing the population along an optimistic-pessimistic coding axis.

### Learning enhances phase-specific communication from the olfactory tubercle to VTA

Having established a learning-dependent temporal reorganization of iDANs, we explored the underlying circuit mechanism by examining simultaneously recorded putative SPNs in the Tu. SPNs exhibited electrophysiological properties characteristic of this cell type, such as low ITI activity, burstiness and monophasic action potentials (Fig. 4A). We found that SPNs mirrored the changes seen in the VTA: while the fraction of task-responsive SPNs increased with learning (Fig. 4B), the overall fraction of coupled units and coupling strength did not change (Fig. 4C). In contrast to iDAN, SPN population activity discriminated stimulus identity even in the untrained animals when looking at units that were coupled to the sniff at the expiration/inspiration-transition during ITI (Fig. S4B, blue). The preferred sniff-coupling phase of SPN also underwent a learning-dependent shift. In expert animals, SPNs developed a strong preference for firing during the post-inspiratory phase, slightly preceding the preferred phase of iDANs (Fig. 4D). With learning, only SPNs coupled to post-inspiration discriminated the stimuli (Fig. S4B, green). In contrast to iDANs, the distributions of preferred coupling phases were bimodal (inspiratory and post-inspiratory) in SPNs both of untrained and trained animals, with the observed shift occurring as a redistribution towards the latter mode. Further characterization of SPN sniff-coupling is provided in Fig. S4C-H.

**Figure 4.**
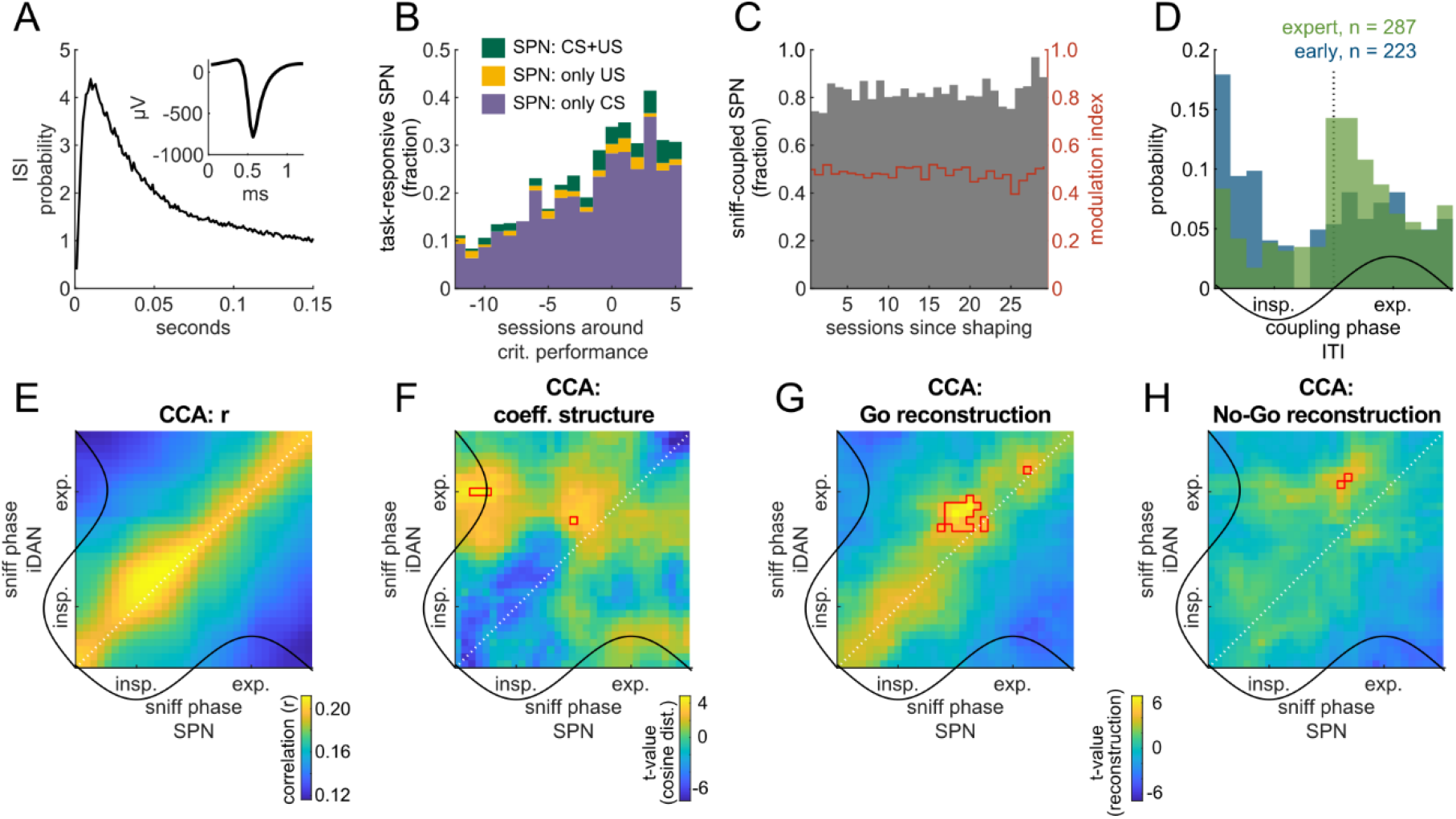
Learning enhances phase-specific Tu→VTA communication. **(A)** Inter-spike interval (ISI) distribution (median) for all task-responsive SPNs, showing characteristic burstiness and long pauses. Inset: Average action potential waveform. **(B)** Fraction of task-responsive SPNs across sessions, aligned to criterion performance. Compared to DANs, US responses are relatively weakly expressed. **(C)** Left axis (gray bars): Fraction of SPNs significantly coupled to the sniff cycle during the ITI across learning sessions. Right axis (red line): Mean modulation index for the sniff coupled population. **(D)** Distribution of preferred ITI coupling phases for task-responsive SPNs in early (blue, n = 223) vs. expert sessions (green, n = 287). The preferred phase shifts significantly to the post-inspiratory/expiratory phase with learning (Watson’s U^2^ test, U^2^ = 0.796, P = 2.98 × 10^−7^), while conserving the initially observed bimodal distribution. **(E–H)** Reconstruction of stimulus responses using phase-resolved functional connectivity models (Canonical Correlation Analysis, CCA) derived from ITI activity. **(E)** Matrix of canonical correlations (R) between SPN and iDAN population activity for each combination of SPN phase and iDAN phase during the ITI in expert animals. **(F)** Matrix showing the structure of the canonical coefficients (measured as the cosine distance to neighboring bins), revealing that neuronal assemblies are more varied (i.e. higher distance) in the post-inspiratory window in Tu→VTA direction. **(G–H)** Reconstruction performance (correlation between stimulus responses of SPN and iDAN projected into the first canonical dimension) for Go trials **(G)** and No-Go trials **(H)** in expert animals. Models derived specifically from the post-inspiratory phase (red outlined cluster) significantly reconstruct the stimulus-evoked activity (Holm-Bonferroni corrected t-test, p < 0.01).

The emerging temporal alignment may arise through an acquired communication channel. To test this, we first constructed functional connectivity models of SPNs and iDANs activity during the ITI with canonical correlation analysis (CCA, see Methods for details). The magnitude of the canonical correlation was high across the phase pairings along the diagonal (i.e. when SPN and iDAN phases were aligned), indicating a general co-modulation of the two neuron types (Fig. 4E). However, this analysis of correlation strength did not identify a privileged phase for communication. Instead, the structure of the neural ensembles defining the correlation (cosine distance between phase-phase-combinations of canonical coefficients) appeared significantly more varied during the post-inspiratory phases (Fig. 4F). The functional relevance of these specific ensembles became clear when we used these ITI-derived models to reconstruct stimulus-evoked responses. Only the connectivity models derived from the post-inspiratory phases with SPNs preceding iDANs significantly reconstructed the conditioned stimulus response in expert animals (Fig. 4G,H), a relationship absent in untrained animals (Fig. S4I-L). This reveals that while correlated activity exists across the sniff cycle, a specific channel for processing task-relevant information is gated during the post-inspiratory phase.

Finally, we probed the behavioral significance of this channel by modeling its strength on a session-by-session basis. Stimulus reconstruction had identified the functionally relevant phase window, yet the raw CCA correlation values largely reflected the prominent zero-lag co-modulation shared by both populations (Fig. 4E, diagonal). To isolate the directed interactions hidden within this global rhythm and measure channel strength and directionality, we employed pairwise cross-correlation analysis (see Methods). With learning, a significant asymmetrical connection emerged, with a peak in SPN activity leading iDAN activity by approximately 20 ms (Fig. 5A, B). Significantly cross-correlated pairs appeared to be coupled more tightly to the post-inspiratory phase (Fig. S5A, B). Crucially, this enhanced communication was phase-specific: the probability of finding the strongest correlation was highest for iDAN spiking during the post-inspiratory phase (Fig. 5C, green). A peak, albeit less pronounced, could already be observed during the shaping session, when gross neural activity had not yet shifted to the post-inspiratory phase (Fig. 5C, blue).

**Figure 5.**
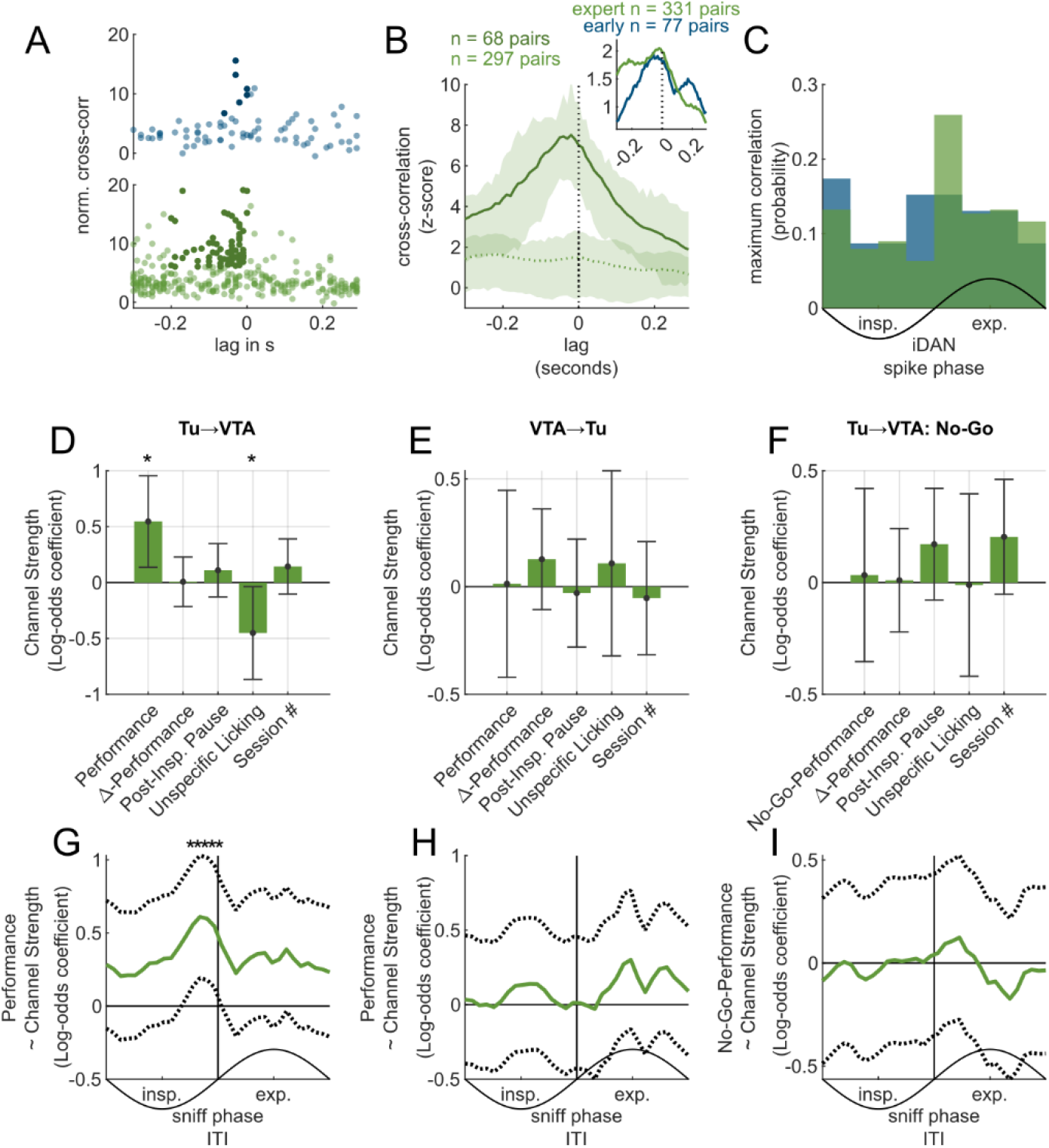
Tu→VTA channel strength tracks task performance during learning. **(A)** Peaks and lags of normalized cross-correlograms for all recorded SPN-iDAN pairs in early (blue, n= 77) and expert sessions (green, n= 331). Each dot represents one pair. Significantly correlated pairs are shown in darker colors. **(B)** A peak emerges at negative lags (∼20 ms with SPNs leading iDANs) in the mean z-scored cross-correlogram for significantly cross-correlated pairs (n = 68 pairs) compared to non-significant pairs (n = 297) in expert animals. Inset: Mean z-scored cross-correlation between SPN-iDAN pairs in early (blue) and expert sessions (green). **(C)** Probability that the maximum cross-correlation for a given pair occurs when the iDAN spike is in a specific phase bin. The post-inspiratory phase contributes most to the correlation in expert (green) vs. early sessions (blue). **(D–F)** Results of a generalized linear mixed-effects model (GLME) predicting session-wise Tu→VTA channel strength from behavioral and state variables. Bars represent log-odds coefficients for each predictor; positive values indicate a positive relationship. Error bars represent the 95% CI. **(D)** Model for discrimination learning sessions (n = 70). Performance is a significant positive predictor. **(E)** Model for VTA→Tu directionality, demonstrating that the performance is linked specific to the Tu→VTA directionality. **(F)** Model for No-Go trial performance, showing no relationship with channel strength. **(G–I)** Phase-resolved analysis of the GLME coefficient for behavioral performance predictors. **(G)** Tu→VTA: The relationship between performance and channel strength is significant only for channel strength derived from the post-inspiratory phase. **(H)** Performance coefficient for VTA→Tu channel strength shows no sniff-phase dependency, as does **(I)** the No-Go Performance coefficient for Tu→VTA.

**Figure 6.**
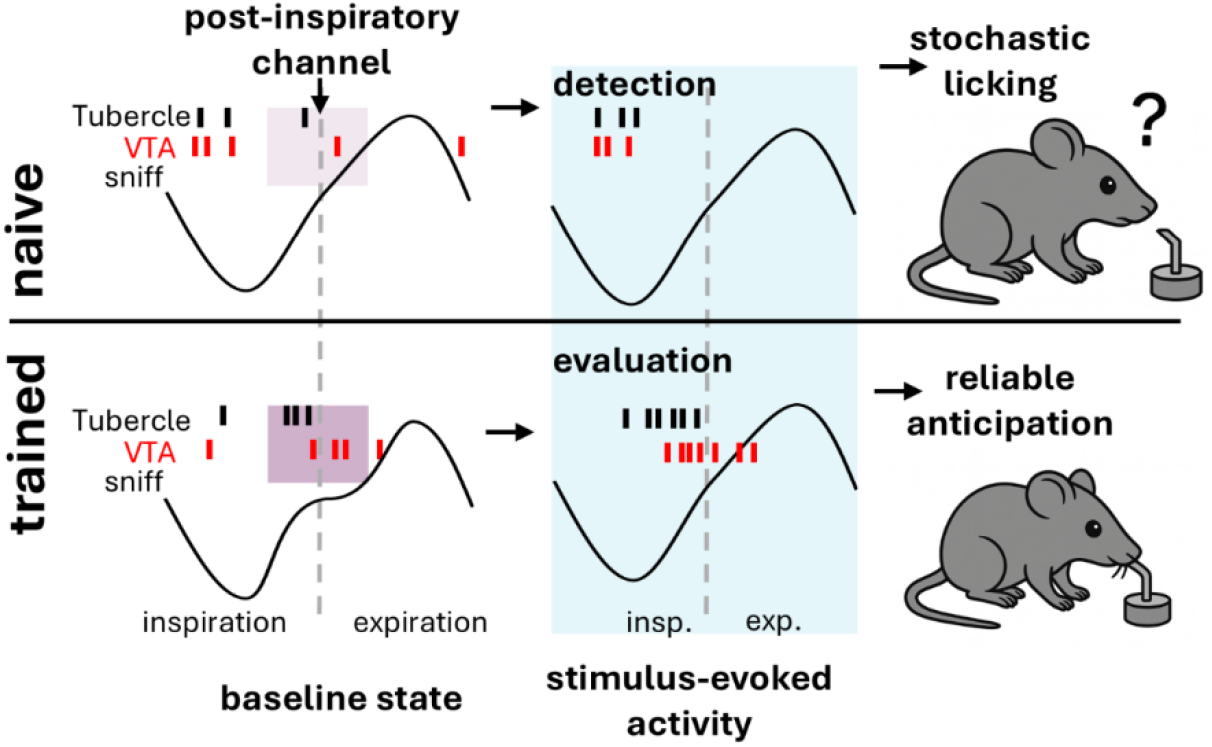
Learning-dependent gating of a Tu→VTA communication channel. The schematic illustrates the proposed functional reorganization of the olfactory tubercle (Tu) to ventral tegmental area (VTA) circuit across two learning states (naïve vs. trained) and two task epochs (baseline state, stimulus-evoked activity). The sniff cycle provides a temporal scaffold for neural activity (black ticks: Tu spiny projection neuron spikes; red ticks: VTA dopamine neuron spikes). *Naive State (Top Row):* In naive animals, baseline Tu and VTA activity is coupled to the inspiration phase of the sniff cycle. A post-inspiratory Tu→VTA channel emerges in the absence of reconfiguration of bulk activity and already reflects emerging anticipatory behavior. During stimulus presentation, responses peak mostly during early inhalation and presumably reflect detection rather than evaluation. *Trained State (Bottom Row):* After learning, the circuit is reconfigured. During the baseline state, Tu and VTA activity becomes organized around the post-inspiratory phase strengthening the communication channel: a temporal window of heightened, phase-locked directional Tu→VTA coordination (visualized by darker shading of the purple box). Concurrently, the post-inspiratory phase lengthens with learning, widening the channel. During stimulus presentation, peak responses occur later and reflect information about the reward probability of the stimuli. This coordinated state thus allows for the discriminative evaluation of the olfactory stimuli, gating the expression of reliable anticipatory behavior.

Using a generalized linear mixed-effects model (GLME) for sessions when the animals’ performance had not yet reached criterion, we found that task performance was a significant positive predictor of channel strength (Fig. 5D). This relationship was sniff-phase-specific, with the model coefficient for performance peaking during post-inspiration (Fig. 5G). Furthermore, the effect was direction-specific and was absent when testing for information flow in the direction VTA-to-Tu direction (Fig. 5E, H). Notably, the relationship in Tu→VTA direction was already present during initial shaping sessions (Fig. S5C, G), a period before the population-level iDAN activity had quantitatively shifted to this phase and before the systematic prolongation of the post-inspiratory phase (cf. Fig. 2). This suggests that the directional functional communication channel is already present in this temporal window prior to the phase-specific consolidation of bulk neural activity.

To delineate specific contributions to this Tu→VTA channel from intercorrelated behavioral and state variables, we included several factors in the model. For the above-described cases (during initial shaping and discrimination learning), the Tu→VTA channel’s relationship with performance was significant (Fig. 5D, Fig. S5C, D) after accounting for the influence of task experience (‘session #’), sniff shape (‘post-insp. pause’), the rate of intra-session learning (‘delta-performance’), and a measure of overall motor activity (‘unspecific licking’). For unspecific licking, we found a negative relationship with channel strength (Fig. 5D, Fig. S5C,D). These results suggest a role for the channel that is specific to task-contingent performance rather than general motor activity or learning rate. Additional analyses revealed the effect was driven by performance on Go-trials (Fig. S5D, H), with no significant relationship between channel strength and No-Go performance during discrimination learning (Fig. 5F, I). This pattern of results is consistent with a single, unified behavioral function: the Tu→VTA channel facilitates the expression of the learned anticipatory behavior associated with olfactory stimuli.

## Discussion

We find here that VTA DAN activity is systematically modulated by the sniff cycle, imposing a continuous temporal structure to their operation. This links dopamine’s role in olfactory learning to brain-wide respiratory entrainment and suggests that this ongoing sensory rhythm is integral to how predictive signals are formed.

Respiration-entrained oscillations are detectable in cortical and subcortical structures and are discussed as a pacemaker for state-dependent coordination (30, 32, 33, 35, 37). A mechanistic framework for how such rhythms could shape information transfer is “communication through coherence” (CTC): rhythmic synchronization generates recurring windows of elevated postsynaptic sensitivity, such that inputs arriving consistently during high-gain phases are transmitted more effectively (19). In this sense, respiration-phase locking can be interpreted as temporal segmentation of the neuronal code. Our findings reveal a role of respiration-phase locking in the mesolimbic system: although Tu and VTA co-vary across the sniff cycle, the task-relevant reconstructive coupling concentrates in a narrow post-inspiratory window, consistent with a phase-selective effective connectivity for interareal communication. The lag observed in our data (Tu leading VTA by ∼20 ms) is compatible with a model in which Tu units are preferentially effective when VTA DANs are in a high-gain post-inspiratory phase, with the strength of this channel tracking task performance. Future closed-loop studies may explore how phase-timed perturbation Tu activity patterns may influence predictive behavior of animals.

The learning-dependent shift in the preferred respiratory coupling phase of DANs suggests that this temporal scaffolding reconfigures with learning. This retuning concentrates the DAN spiking within the post-inspiratory phase of the sniff cycle. A neuron’s inter-trial-interval coupling phase predicts its subsequent participation in encoding the conditioned stimulus, consistent with a phase-dependent state of ‘readiness’ to engage predictive olfactory circuits. Over the course of learning, SPNs initially code mainly for stimulus identity and, with learning, concentrate their responses in the post-inspiration phase. The bimodal phase distribution of preferential inspiratory versus post-inspiratory firing in SPN upon learning suggests that at least two functionally distinct Tu populations. A plausible but speculative interpretation is that this bimodality partially reflects known functional differences between D1- and D2-expressing SPNs. Imaging in the olfactory tubercle indicates, for example, that D1 neurons can represent odor valence relatively independently of chemical identity, whereas D2 representations can be weaker, more behavior-contingent, and more identity-selective (38). This division also aligns with recent circuit-level accounts in which opponent D1/D2 pathways preferentially represent different “tails” of a reward distribution, thereby supporting distributional reinforcement learning computations (17). This interpretation remains tentative because we did not perform cell-type-specific identification in the present dataset; direct tests will require a better understanding of the interactions of D1- and D2-type receptor-mediated plasticity rules in vivo SPNs and their systematic differences in preferred sniff phase coupling (39, 40). Against this background, phase-dependent segmentation across the sniff cycle provides an additional axis for separating identity-related processing in inspiration-coupled SPN from outcome prediction in SPN preferentially coupled to the post-inspiratory phase.

During the stimulus response itself, information content changes within a single sniff. In naïve animals, activity is concentrated in the early inhalation phase, as expected from bottom-up processing with odor detection or salience (41–43). In contrast, value-related signals emerge later in the sniff cycle. The observed intra-sniff progression from earlier activity consistent with detection/salience to later activity more consistent with value-related signals fits the idea that a single respiratory cycle can function as an elementary time unit for perception and action-guiding evaluation (43, 44). The shaping of the Tu→VTA communication comes with the learning-related shift of dopaminergic and striatal evaluation components toward the post-inspiratory phase. As such, respiration could serve as a carrier in a time-division multiplexing of information in DANs.

Moreover, a DAN’s coupling phase after post-inspiration predicts its position along an optimism–pessimism axis of predicted outcomes derived from distributional reinforcement-learning frameworks. The neurobiological relevance of these frameworks is supported by prior evidence that DANs’ heterogeneous cue responses tile expectiles of future reward so that the population represents a return distribution rather than a single mean (45). Recent work further suggests that striatal D1 vs. D2 pathways may provide a structural basis for preferential representation of different distributional flanks (17). Our findings add a dynamic dimension to this picture: if sniff phase systematically covaries with a dopamine neuron’s position along an optimism–pessimism axis, the respiratory cycle could function as a temporal addressing system that organizes distributional predictions not only across cell types/populations but also via phase-dependent routing. Here, the interaction preferred firing phase and dopamine-mediated spike timing dependent plasticity rules may interact in non-linear fashion (40) to generate such distributional coding of predicted outcomes.

We find that sniffing behavior co-evolves with neural timing: the emergence of a prolonged post-inspiratory pause coincides with a shift of DAN coupling to that same pause but does not determine it, suggesting a synergy between sensing and neural processing. Olfaction is a prototypical active-sensing system: a sniff is (analogous to a saccade or whisking) a unit of active sampling that is modulated by state, task, and expectation (44). In freely moving paradigms, mice couple sniffing to movement and investigation motifs (46). Complementarily, experimental work has shown that dopaminergic signaling can drive changes in sniffing (36) and a prolonged post-inspiratory pause has been linked to reward anticipation in freely moving mice (46). Taken together, the learning-related extension of the post-inspiratory pause could be interpreted as a functional behavioral adaptation that expands the temporal window for phase-dependent Tu→VTA integration. Furthering our findings in a freely moving paradigm may help better understand these mutual interactions.

In sum, the rhythm of sniffing provides a temporal substrate for the mesolimbic dopamine system. Learning reorganizes coupling toward post-inspiration and strengthens Tu–VTA connectivity, imposing a critical dimension of temporal structure on dopamine computations. This offers a neurophysiological instantiation of Communication-Through-Coherence mediated by a sensory rhythm, advancing a mechanistic account of how the brain implements reinforcement learning.

## Methods

### Ethics statement

All procedures were in accordance with the National Institutes of Health Guide for the Care and Use of Laboratory Animals and the EU 2010/63 directive, and approved by the local animal welfare authority (Referat 35, Regierungspräsidium Karlsruhe, Karlsruhe, Germany).

### Subjects

Subjects were 8 adult male and female DAT:(IRES)Cre^+^ mice (B6.SJL-Slc6a^3tm1.1(cre)Bkmn^/J; RRID: IMSR_JAX:006660, Jackson Laboratory) (47) crossed with a Cre-dependent Channelrhodopsin-2 reporter line (Ai32(RCL-ChR2(H134R)/EYFP, B6;129S-Gt(ROSA)^26Sortm32(CAG-COP4*H134R/EYFP)Hze^/J, RRID:IMSR_JAX:012569, Jackson Laboratory) (48) resulting in dopamine neuron-specific expression of ChR2 (Fig. S1C). All animals had been kept in a C57BL/6J (Charles River, Sulzfeld, Germany) for F>10. Animals were housed on a 12-hour light/dark cycle and had ad libitum access to food. Water access was restricted prior to behavioral training sessions with daily wellness checks and weighing to ensure >85% baseline bodyweight.

### Surgical Procedures

A total of 10 mice were implanted with custom-built, fixed dual-region microelectrode arrays designed for chronic recordings from the olfactory tubercle (Tu) and ventral tegmental area (VTA). Two animals were excluded from analysis due to insufficient numbers of dopamine neurons. Each array consisted of 16 tetrodes per target region, spun from 12.5 µm tungsten wire (California Fine Wire). Tetrode tips were gold-plated to a target impedance of ∼100 kΩ using a NanoZ device (Multi Channel Systems). Each array also included one optical fiber (200 µm, 0.39 NA, Thorlabs) per region, positioned above the tetrodes for optogenetic stimulation.

For surgery, animals received analgesic (meloxicam, Metacam Boehringer Ingelheim) before and after surgery. Mice were anesthetized with isoflurane (1-2% in pure O_2_), with the concentration titrated to maintain a stable breathing rate of approximately 1 Hz. Local anesthetic (Xylocaine) was applied to the scalp prior to incision. The animal was then fixed in a stereotactic frame (Kopf Instruments). After the scalp was incised and retracted, the wound margins were attached using 3M VetBond tissue adhesive, ensuring temporal and occipital muscle insertions were not damaged. Two craniotomies were made over the target locations (relative to bregma, midpoint of Tu: 1.7 mm ML (left hemisphere), 1.5 mm AP, 5.5 mm depth; VTA: 0.5 mm ML (bilateral), −3.5 mm AP, 4.5 mm depth) with a pneumatic drill, and the dura mater was resected. Ground and reference gold pins (Neuralynx) were implanted over the cerebellum. The dual-region array was lowered to its target coordinates using an automated micromanipulator (Luigs & Neumann), with the final descent occurring over a 15-minute period to minimize tissue damage. Once in position, the array was secured to the skull using dental cement (C&B Superbond, Sun medical for adhesion to skull, Kulzer Palladur to encase the implant). A 3D-printed titanium headbar was fixed to the implants. The final weight of the implant, including cement, was less than 3 g.

### Histological Verification of Recording Sites

Histological verification of electrode placement was performed using a rapid, stain-free imaging method based on intrinsic tissue contrast (49). Following the final recording session, animals were deeply anesthetized with ketamine/xylazine (300 mg/kg BW ketamine and 60 mg/kg BW xylazine diluted in 0,9% saline, i.p. injection) and transcardially perfused with phosphate-buffered saline followed by 4% paraformaldehyde in saline. Brains were extracted, post-fixed in 4% paraformaldehyde, and embedded in agar for stability during sectioning. Sagittal sections (100 µm) were cut on a microtome and transferred directly into a dish containing cold phosphate-buffered saline. Sections were then mounted on glass slides with buffer and immediately imaged in their wet state using a brightfield microscope (Neurolucida). This provided intrinsic optical contrast by differential light scattering between the mechanically disrupted tissue of the electrode track and the surrounding, more uniform neuropil, while the chosen thickness allowed easy tracking over multiple sections. By imaging in a hydrated state, this method minimizes the spatial distortion and shrinkage artifacts associated with dehydration and mounting procedures. Clear visibility of electrode traces allowed confirmation of placement in the targeted areas.

### Olfactometer

All behavioral training and recording sessions were conducted in a custom-built sound-insulated faraday chamber. Head-fixed mice were positioned facing a custom-made face mask that served as the odor delivery port with an integrated pressure sensor (HDIMOLOGBY8H3, Sensor Technics). Licking was detected by an infrared beam-break sensor (SX-3009-P1, Omron Electronics) integrated into the face mask in front of the lick spout. Water rewards (∼3 µL) were delivered to the lick spout via a gravity-fed system controlled by a solenoid valve (NResearch).

Odors were delivered using a custom-built olfactometer. Pure odorants (Ylang-Ylang, Geranium, Neroli, Eugenol; all from Sigma-Aldrich) were diluted in mineral oil to 0.5% concentration. A constant stream of filtered air was passed through mass flow controllers (Natec Sensors) to generate a constant clean air carrier stream. During stimulus presentation, solenoid valves (NResearch) controlled by a micro-controller (Arduino Mega) diverted the air stream through one of the odorant vials to pre-flush PTFE tubing and equilibrize pressure. Odor delivery was then effected by a final valve, switching between clean air and odorized air to the animal’s face mask. Both pre-loading and odor delivery tubes were switched to clean air at odor offset. The timing of odor and reward delivery was controlled by custom MATLAB scripts.

### Behavioral Task and Training Stages

Mice were trained on a head-fixed, olfactory Go/No-Go discrimination task. Each trial began with a variable inter-trial interval (ITI) of 8-12 seconds, during which the animal received only clean air. This was followed by a 1-second odor presentation (conditioned stimulus, CS). A 1-second response window began at the offset of the CS, during which the licking was considered as anticipatory. The reward was delivered at the end of this response window.

### Training proceeded as follows

#### Shaping

Naive animals were first trained to associate a single odor (CS) with a 100% water reward. Animals learned to lick after stimulus presentation to obtain the reward.

#### Discrimination

Once proficient in the shaping task, animals were introduced to the full three-odor task. Three distinct odors were presented in a pseudorandom order, associated with different reward probabilities: a high-value odor with 100% reward probability, a medium-value odor with 50% reward probability, and a low-value odor with 0% reward probability. Trials with 100% and 50% reward probability were considered “Go” trials, where licking was the correct action. Trials with 0% reward probability were “No-Go” trials, where withholding licks was the correct action.

For both stages, session performance was calculated as the fraction of correct trials. A trial was considered correct if the animal licked during the response window on a Go trial (a “Hit”) or correctly withheld licking for the entire response window on a No-Go trial (a “Correct Rejection”).

### Data Acquisition

All electrophysiological and behavioral data were acquired using an Intan Technologies recording system. Wideband neural signals were recorded from two 64-channel RHD headstages (Part #C3315) connected to a USB Interface Board (Intan RHD Recording Controller). The signals were digitized at the headstage at a rate of 30 kS/s and streamed to the acquisition board. Behavioral and task-related events were recorded concurrently and synchronized with the neural data stream via the acquisition board’s digital and analog inputs. The onset and offset of odor delivery, licking, laser stimulation for optogenetic tagging, and reward delivery were recorded as digital TTL inputs. The respiratory signal (sniffing) was recorded as a continuous analog input from the pressure sensor embedded in the animal’s face mask.

### Spike Sorting and Unit Curation

Spike sorting was performed using Kilosort 3. The output of the initial sort was then processed through an automated post-hoc curation pipeline to address misclassifications and ensure high unit quality. This pipeline applied the following sequential criteria:

#### Spatial Specificity

For each unit, the mean squared energy of its average waveform on its primary tetrode was compared to the energy on adjacent tetrodes. Units with an energy ratio below 10 were considered to have waveforms that were not spatially localized to a single tetrode and were excluded.

#### Refractory Period Violations

The fraction of inter-spike intervals shorter than 2 ms was calculated for each unit. Units with a violation rate exceeding 2% were excluded.

#### Lick Artifact Removal

To identify potential contamination from licking-related electrical artifacts, the temporal overlap between spike times and lick times was assessed. Units with spike times that coincided with more than 90% lick events were classified as artifacts and excluded.

#### Redundant Unit Merging

To correct for instances where a single neuron was split into multiple clusters because of action potentials with long duration and high amplitude (which concerned dopamine neurons), we identified pairs of units recorded on the same tetrode with highly overlapping spike trains (>70% of spikes from one unit occurring within 1 ms of a spike from the other). In such cases, the unit with fewer spikes was merged into the unit with more spikes. This process was applied iteratively to resolve sequential and circular merge chains.

Units that passed all criteria were retained for subsequent analyses.

### Optogenetic Tagging for iDANs and classification of SPNs

To identify iDANs, we ran an optogenetic tagging protocol at the end of each recording session. A train of 200 light pulses (10 Hz, 473 nm, pulse width 5 ms, power approximately 5 mW at fiber tip) was delivered through the optical fiber targeting the VTA.

For each recorded VTA neuron, a spike-laser cross-correlogram was computed using a 1 ms bin size over a ±20 ms window around each laser pulse onset. To statistically assess the significance of any short-latency activation, a null distribution was generated using a pulse dithering method. For each of 1000 iterations, a random temporal dither (uniformly distributed between 0 and 60 ms) was added to each laser pulse time, and a control cross-correlogram was computed. A global 97.5th percentile confidence limit was calculated from the distribution of all bins across all 1000 control correlograms. A neuron was classified as an iDAN if its real cross-correlogram showed significant activation, defined as at least two consecutive 1-ms bins within the 20 ms post-stimulus window exceeding this 97.5th percentile threshold.

SPNs were classified as all recorded units in the olfactory tubercle below a 5 Hz ITI firing rate.

### Sniff Phase Extraction and Post-Inspiratory Pause Quantification

The raw respiratory pressure signal, originally sampled at 100 Hz, was upsampled to 500 Hz using interpolation and then high-pass filtered at 0.2 Hz using a 3rd-order Butterworth filter to remove slow drifts. To normalize changes in sniff amplitude, the signal was divided by its envelope, which was calculated using the envelope function in MATLAB. The instantaneous sniff phase was then extracted from this normalized signal using the Hilbert transform. The resulting phase angle was such that the beginning of inspiration corresponds to a phase of −π, and the beginning of expiration corresponds to 0.

To quantify the duration of the post-inspiratory pause for each sniff, we analyzed the recorded pressure between the points of maximum inspiration and maximum expiration. The waveform in this window was normalized to a range of [0, 1]. The first and second derivatives of this normalized waveform were computed. The start of the pause was defined as the first prominent local minimum in the second derivative following the initial peak in the first derivative. The end of the pause was defined as the subsequent maximum peak in the second derivative. For a visualization of the method, see Fig. S6. The duration of the pause was calculated as the time between these two points, normalized by the total duration of the sniff cycle.

### Analysis of Sniff-Phase Coupling

To quantify the coupling of neural firing to the sniff cycle, we performed an analysis based on the instantaneous firing rate of each neuron. First, a continuous spike density function was generated for each neuron by convolving its spike train with a 150 ms Gaussian kernel.

For a given behavioral epoch (e.g., the inter-trial interval), all individual sniff cycles were identified from the respiratory pressure signal. For each of these sniff cycles, the corresponding segment of the neuron’s spike density function was sampled into 33 phase bins, spanning from −π to π, according to the instantaneous sniff phase. This approach is robust to the non-sinusoidal nature of the sniff waveform (i.e. the presence of post-inspiratory pauses), as it ensures that the firing rate is correctly mapped to the corresponding phase regardless of temporal distortions within the cycle. This procedure resulted in a matrix where each row represented a single sniff cycle and each column represented a phase bin, with the values being the instantaneous firing rate.

A neuron was then classified as significantly coupled to the sniff cycle if the distribution of its instantaneous firing rates at its preferred (peak) phase was significantly different from the distribution at its anti-preferred (trough) phase across all sniff cycles in the epoch (Wilcoxon rank-sum test, p < 0.05). The preferred coupling phase for each neuron was defined as the center of the phase bin with the highest mean instantaneous firing rate. A modulation index was calculated from this mean phase-rate histogram as (rate_max_ - rate_min_) / (rate_max_ + rate_min_).

### Analysis of Stimulus Discriminability by ITI Coupling Phase

To test whether a neuron’s preferred sniff-coupling phase during the inter-trial interval (ITI) related to its ability to encode stimulus value, we performed a sliding window analysis. First, for each neuron, we quantified its stimulus discriminability as the chi-squared statistic from a Kruskal-Wallis test comparing its responses across the three odor conditions. We then assessed how this discriminability metric was distributed across the population as a function of preferred ITI phase. A circular window (width: 1.5 rad) was moved across the phase range from −π to π in steps of 1/3 rad. For each window position, we identified all neurons for which preferred ITI coupling phase fell within the window’s bounds. For each resulting group of neurons, we calculated the mean discriminability score. To determine the statistical significance of the mean value in each window, we performed a permutation test. A null distribution was generated by randomly shuffling the discriminability scores among all neurons and recalculating the mean for the neurons within the window (1000 iterations). The p-value was calculated as the proportion of shuffled means that were greater than or equal to the true observed mean. Bins with p < 0.05 were considered significant. To visualize the uncertainty of the mean, a 95% confidence interval was also calculated for each window using a bootstrap procedure (1000 iterations).

### Trial-by-trial CS Response Modulation by ITI Phase

To test if a neuron’s stimulus response on a given trial was modulated by its sniff-coupling state in the immediately preceding inter-trial interval (ITI), we performed a sliding window analysis at the single-neuron level. First, for each neuron, we determined the preferred ITI sniff phase for every trial by finding the peak of the interpolated phase-rate histogram for that trial. The neuron’s corresponding stimulus response on each trial was z-scored relative to all other trials for that neuron. This resulted in a set of (phase, z-scored response) pairs for each neuron. Next, a continuous phase-response trace was calculated for each neuron. A circular window (width: 1.5 rad) was moved across the phase range from −π to π in steps of 0.33 rad. The value of the trace at each window center was the mean z-scored response of all trials from that neuron whose ITI phase fell within the window. To ensure robust analysis, only neurons that had trials in every window position were included for subsequent population analysis. The mean trace was then calculated across this population of neurons. To assess the significance of the observed modulation, a permutation test was performed. For each of 1000 iterations, the phase-response relationship was broken by shuffling the binned window values within each neuron independently. A null distribution of mean population traces was generated from this procedure. The final trace was visualized by z-scoring the real mean trace relative to the mean and standard deviation of this null distribution at each phase bin. Significance thresholds (p < 0.05) were calculated as the 2.5th and 97.5th percentiles of the null distribution. The certainty of the mean trace estimation was visualized by a 95% confidence interval, calculated by bootstrapping (resampling neurons with replacement, 1000 iterations) and normalizing the resulting traces in the same manner. An overall p-value for the modulation depth was calculated by comparing the range (max-min) of the real mean trace to a null distribution of ranges from the permuted traces.

### Distribution Decoding

We replicated the decoding analysis by Dabney et al. using their published Python code (16). In short, we quantified the mean normalized response to all three CS for each neuron, yielding the response matrix V. Asymmetric scaling factors τ were approximated via the relative position of the normalized CS_50%_ response within the population (Fig. S3D). Decoding consists of minimizing via a truncated Newton method (SciPy 1.2.1) the loss given by

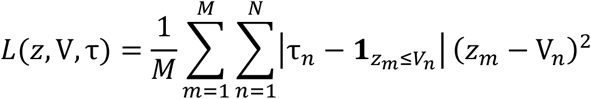

with *z* being one of M=100 random samples, N being the number of neurons and **1** being the indicator function. As described in Dabney et al., the minimization was initiated at the sample points with the smallest loss of 20000 random samples uniform in the range of rewards. The line plot in Fig. 3H shows the resulting samples smoothed with a kernel density estimation.

### Value Encoding by Peak Response Phase

To analyze the temporal evolution of value coding within a single sniff, we first calculated a phase-resolved stimulus response for each neuron. For each of the 33 sniff phase bins, we computed the auROC for the high, medium, and low reward odors relative to a pre-stimulus baseline.

For each neuron, we then identified its ‘peak response phase’ as the single phase bin in which it showed the most significant discrimination between the three odor conditions (i.e., the bin with the minimum p-value from a Kruskal-Wallis test on the response distributions).

At this specific peak response phase, we calculated a normalized metric for the coding of the 50% reward odor, referred to as “value of 50% odor” in Fig. 3I. This was calculated as (auROC_50%_ – auROC_0%_) / (auROC_100%_-auROC_0%_), which places the response to the 50% odor on a scale from 0 (encoded like the 0% odor) to 1 (encoded like the 100% odor), discarding units that did not obey the ordering. The circ stat toolbox was used to compute circular-circular and circular-linear correlations (50).

### Cross-Correlation and Functional Connectivity Analysis

To quantify the functional connectivity between Tu SPNs and VTA iDANs, we computed a z-scored cross-covariance of their instantaneous firing rates (IFRs).

First, continuous IFRs for all neurons were calculated by convolving their spike trains with a 150 ms Gaussian kernel, and the resulting signals were downsampled to 100 Hz. For each SPN-iDAN pair, we calculated a time-lagged cross-covariance of their IFRs during the inter-trial interval (ITI). This was done by computing the sum of the cross-products of the two IFR vectors across all ITI trials for time lags ranging from −300 ms to +290 ms in 10 ms steps.

To assess the statistical significance of this raw cross-covariance and normalize for chance correlations, a null distribution was generated via a trial-shuffling procedure. For each neuron pair and each time lag, we created a distribution of 50 surrogate cross-covariance values by randomly shuffling the trial identities of one neuron’s IFR vector before re-computing the cross-product sum. The real cross-covariance value for each time lag was then z-scored relative to the mean and standard deviation of its corresponding shuffled distribution. This procedure yields a z-scored cross-correlogram for each neuron pair, where the value at each time lag represents the strength of the functional connectivity in standard deviations from chance.

### Correlation Probability by Spike Phase

To determine if the strength of the Tu→VTA functional connectivity was modulated by the sniff cycle, we performed a phase-resolved correlation analysis. For each SPN-iDAN pair, we first identified the single time lag at which their cross-covariance was maximal. Then, we identified all time points during the ITI that corresponded to one of the 33 sniff phase bins. At these specific time points, we calculated the Pearson correlation between the IFR of the iDAN and the IFR of the SPN at the pre-determined optimal time lag. This procedure resulted in a vector of 33 correlation coefficients for each neuron pair, describing how the strength of their functional connectivity varied across the sniff cycle. To generate the population summary shown in Figure 4i, we identified the single sniff phase bin in which this correlation was maximal for each neuron pair and constructed a histogram from the resulting distribution of ‘best phases’.

### Phase-Resolved Canonical Correlation Analysis (CCA) for Stimulus Reconstruction

To test if the phase-specific functional connectivity observed during baseline was relevant for stimulus processing, we used a cross-prediction approach based on Canonical Correlation Analysis (CCA). For each session, we first constructed two population data matrices, X (SPNs) and Y (iDANs), from the instantaneous firing rates during the inter-trial interval (ITI). The data was arranged such that the matrices had dimensions of (sniffs * trials) x phase bins, effectively treating each sniff as an independent sample.

We then performed a phase-resolved CCA. For each of the 1089 possible combinations of SPN phase bin (i) and iDAN phase bin (j), we fit a CCA model between the corresponding population vectors X(:,i) and Y(:,j). This procedure yielded a 33×33 matrix of canonical correlations (r) and a set of canonical coefficients (projection vectors A and B) for each phase-pair combination. These coefficients define the specific weighted combination of neurons that form the most correlated dimension between the two populations for that specific phase-pair.

In the final step, we tested if these baseline-derived connectivity models could reconstruct stimulus-evoked activity. We extracted the stimulus-evoked firing rate vectors for SPNs and iDANs during the first sniff after stimulus onset. For each of the 1089 baseline-derived CCA models, we projected the stimulus-evoked SPN and iDAN activity onto their respective canonical dimensions using the coefficients (A and B) derived from ITI. We then calculated the Pearson correlation between these projected time series. This resulted in a 33×33 matrix of reconstruction correlations for each session. To assess significance at the group level, these correlation values were converted to t-statistics, and a one-sample t-map was assembled across sessions. The significance in this t-map was assessed with a Holm-Bonferroni correction for multiple comparisons (p < 0.01).

### Statistical Modeling of Session-Level Data

To investigate the relationship between Tu→VTA functional connectivity and behavior, we performed a session-level analysis using generalized linear mixed-effects models (GLMEs). All analyses were conducted in MATLAB using the fitglme function from the Statistics and Machine Learning Toolbox.

First, a data table was constructed where each row represented a single experimental session. For each session, we calculated several behavioral and state variables. Task performance (Performance) was computed as described above. Intra-session learning (Δ-Performance) was defined as the change in performance between the first and last thirds of the session. A proxy for general motor activity (Unspecific Licking) was calculated as the overall frequency of licking throughout the session. The sniffing behavior (Post-Insp. Pause) was defined as the session-averaged duration of the post-inspiratory pause. Overall task experience was represented by the session number (Session #).

The primary neural variable, Tu→VTA Channel Strength, was derived from the cross-covariance analysis described above. For each session, we first identified the Tu→VTA neuron pair with the maximal cross-covariance at negative time lags (Tu leading VTA), or for the control shown in Fig. S5, at positive time lags (VTA leading Tu). For the initial model, the Channel Strength for that session was then defined as the session-level cross-covariance value (see above) of the strongest interacting pair at the time bin corresponding to the post-inspiratory phase. To account for idiosyncratic differences in signal magnitude between animals, this Channel Strength metric was z-scored within each animal before being entered into the model. For the phase-resolved analysis, models were computed separately with the channel strength derived from each sniff-phase bin.

## Acknowledgements

We are grateful to the current and former members of the Kelsch Lab for fruitful discussions.

