## Supplemental Information for "Sniffing Shapes Dopamine Signals for Reward Prediction"

Max F. Scheller[1,2], Catarina Luís[2], Wolfgang Kelsch[1,2]
[1]Johannes-Gutenberg-Universität, 55122 Mainz, Germany, [2]Zentralinstitut für Seelische Gesundheit, Heidelberg University, 68159 Mannheim, Germany

**This PDF file includes:**

Figures S1 to S6
Table S1

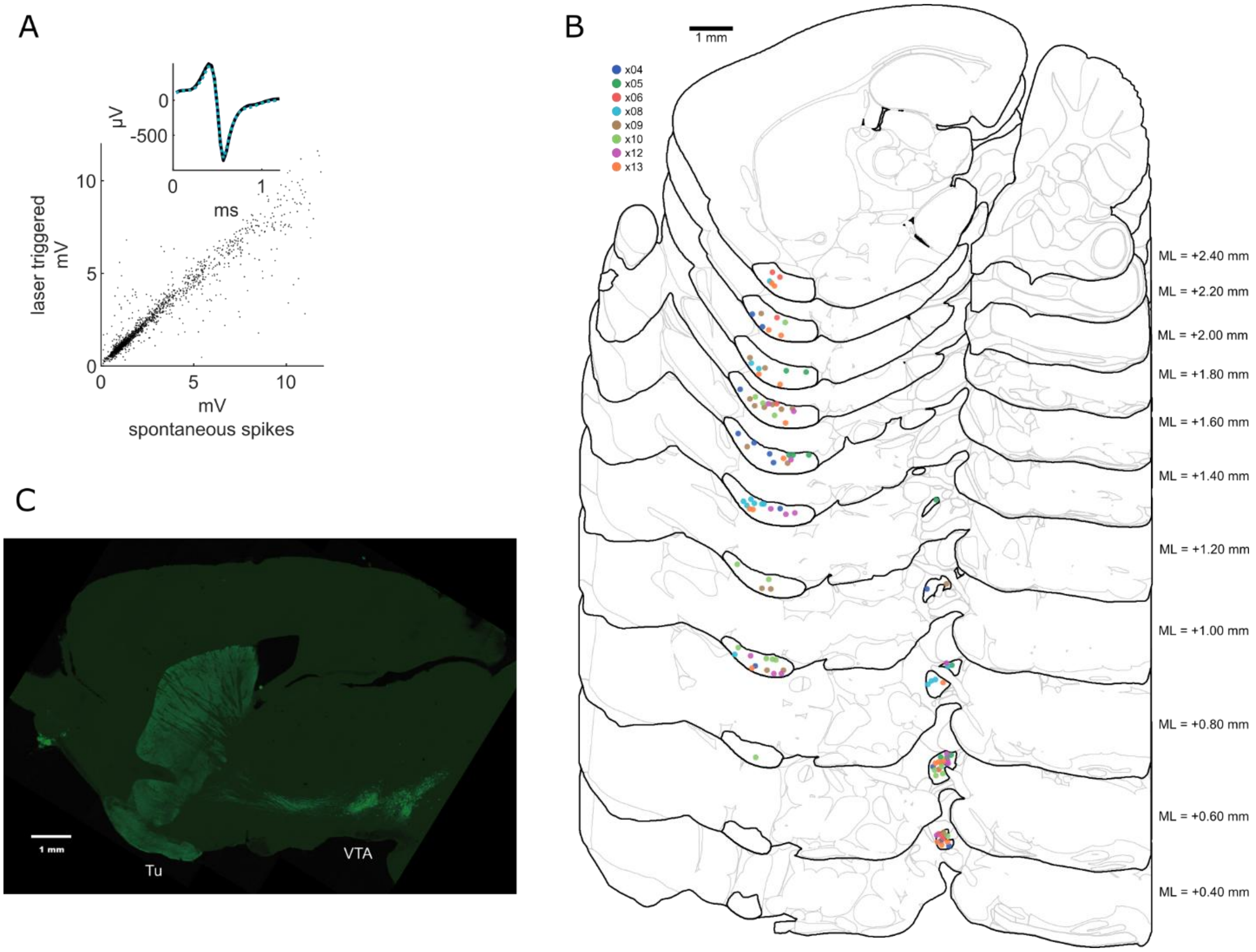


**Figure S1. Validation of optogenetic identification.**

**(A)** The main plot shows a scatter plot of spike amplitudes for all iDANs, comparing the amplitude of laser-triggered spikes versus all other spikes. The linear relationship indicates isolation quality. *Inset:* Overlay of the mean action potential waveform for all iDANs (black) and the mean laser-triggered waveform (blue dashed line), confirming waveform stability during photostimulation.

**(B)** Reconstructed locations of recording tetrodes mapped onto serial sagittal sections for all mice. Circles indicate the recording coordinates for each animal (color-coded). Contralateral VTA tetrode locations are mirrored to the displayed hemisphere for visualization.

**(C)** Pseudocolor image montage showing expression of ChR2-YFP (green) in the VTA and in dopaminergic axons in the striatum and olfactory tubercle (Tu) , scale bar 1 mm.

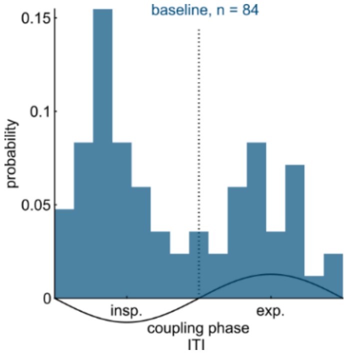


**Figure S2. Sniff coupling of iDANs during habituation.**
Distribution of preferred sniff-coupling phases for iDANs recorded during habituation sessions ($n$ = 84), prior to the start of behavioral shaping. The distribution is bimodal but shows a prominent bias toward the inhalation phase, similar to the early learning phase in Fig. 2D.

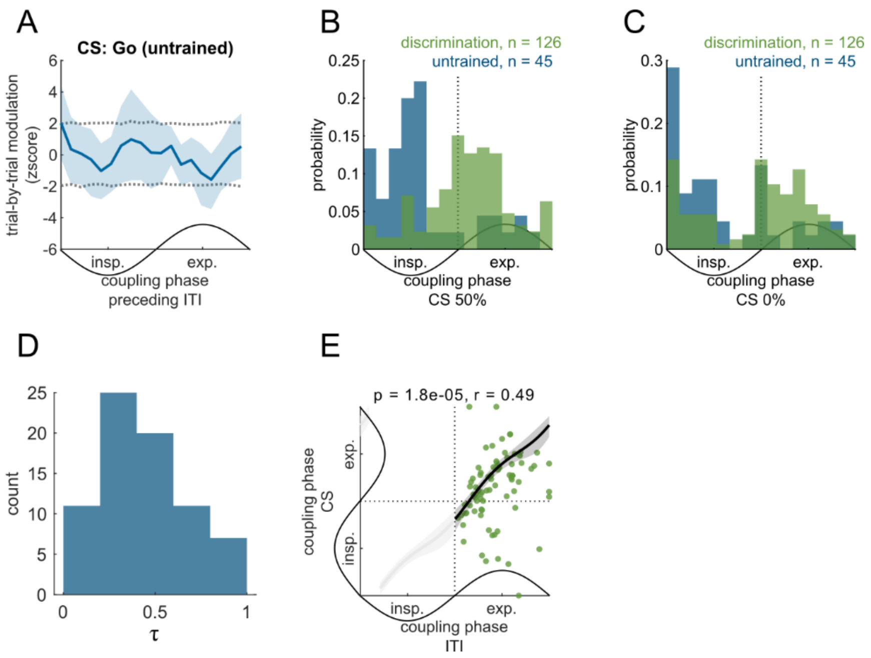


**Figure S3. Sniff-phase coupling and stimulus encoding in untrained animals.**

**(A)** Trial-by-trial modulation of Go-trial CS responses by the preceding ITI coupling phase in untrained animals. Unlike in trained animals (Fig. 3C), there is no significant modulation of the response gain by the phase of the previous sniff.

**(B–C)** Distribution of preferred coupling phases during the CS response for the 50% reward odor **(B)** and the 0% reward odor **(C)** in iDANs from early (blue, $n = 45$) vs. expert (green, $n = 126$) sessions. Similar to the 100% CS (Fig. 3E), the response phase shifts toward expiration in expert animals.

**(D)** Distribution of asymmetric scaling factors τ for the analysis presented in Fig. 3H.

**(E)** Correlation between the preferred coupling phase during the ITI and during stimulus presentation for iDANs (circular-circular correlation, $r = 0.49$, $P = 1.8 \times 10^{-5}$, $n = 80$).

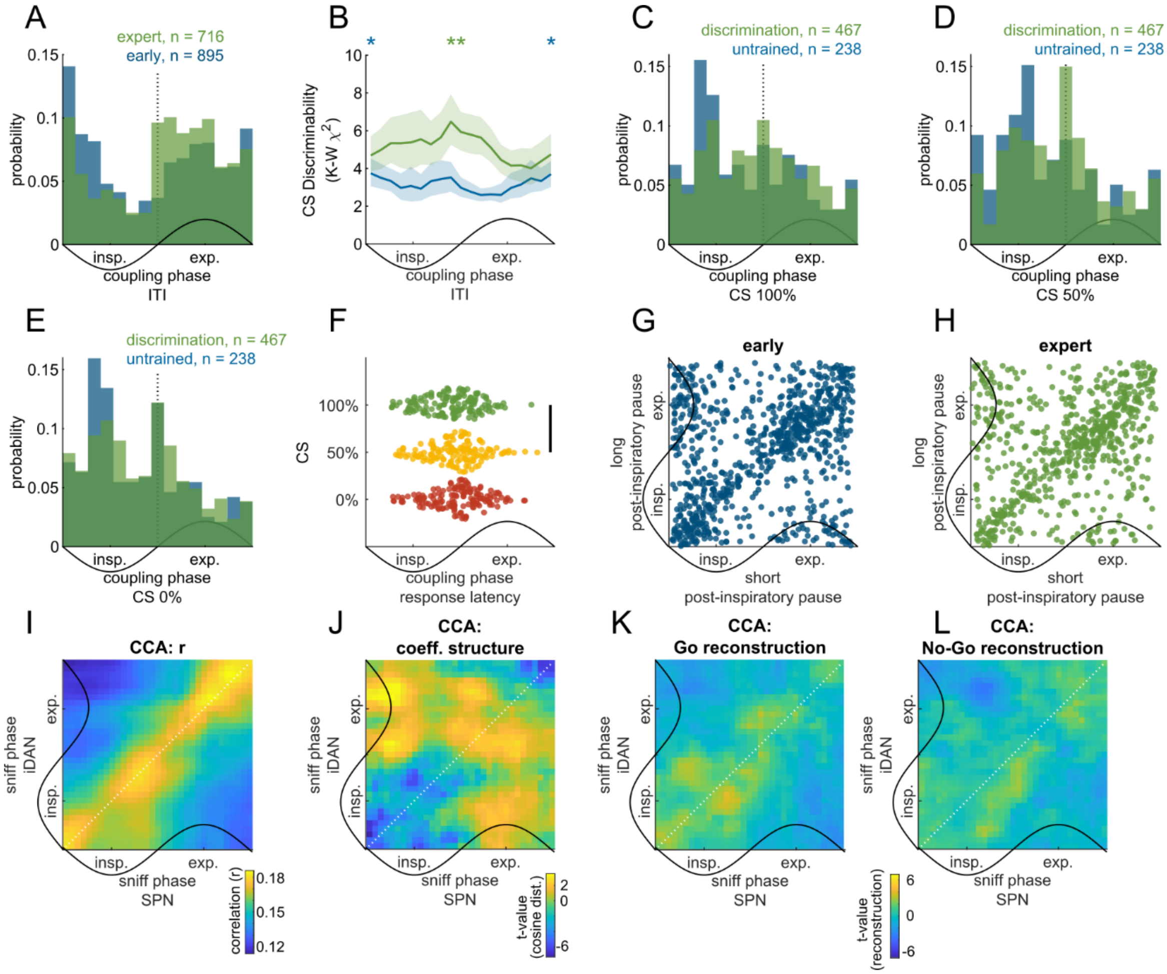


**Figure S4. SPN coupling and connectivity control analyses.**

**(A)** Distribution of preferred ITI coupling phases for all recorded SPNs in early (blue, $n$ = 895) vs. expert (green, $n$ = 716) sessions.

**(B)** SPN discriminability of CS as a function of the preferred ITI coupling phase. In contrast to iDANs, SPNs can discriminate between odors in early training, though this encoding is not aligned to the post-inspiratory phase (identity coding).

**(C–E)** Distribution of preferred coupling phases during the CS response for the 100% **(C)**, 50% **(D)**, and 0% **(E)** reward odors in early (blue) vs. expert (green) sessions.

**(F)** Latency of the first spike burst during the CS presentation for the three reward contingencies.

**(G–H)** Preferred SPN coupling phase plotted against the duration of the post-inspiratory pause for individual sniffs, shown separately for early **(G)** and expert **(H)** training phases.

**(I–L)** CCA-based stimulus reconstruction results for untrained animals (corresponding to the expert analysis in Fig. 4E–H). **(I)** Canonical correlations, **(J)** coefficient structure, **(K)** Go reconstruction, **(L)** No-Go reconstruction. No significant stimulus reconstruction is observed in the untrained state.

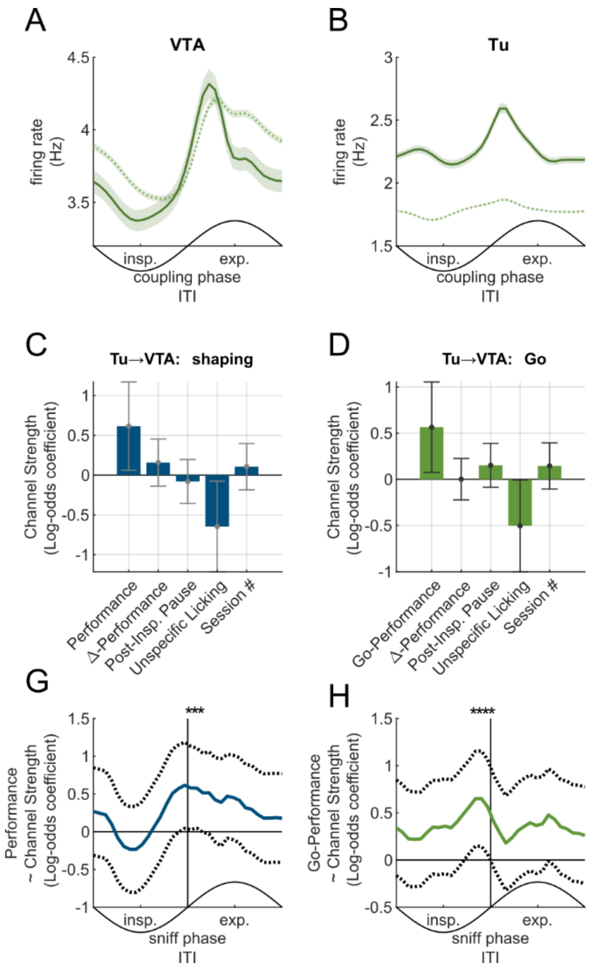


**Figure S5. Control analyses delineating channel function.**

**(A–B)** Mean firing rate as a function of sniff phase for iDANs **(A)** and SPNs **(B)**. Solid lines represent neurons that were part of significantly correlated pairs; dashed lines represent non-correlated pairs. Correlated neurons show tighter phase-locking to the sniff cycle.

**(C–D)** GLME results predicting Tu→VTA channel strength during initial shaping sessions **(C)** and for Go-trial performance during discrimination sessions **(D)**. Performance remains a significant positive predictor even in these specific subsets.

**(G–H)** Phase-resolved GLME coefficient for the *Performance* predictor during shaping **(G)** and for *Go-Performance* during discrimination **(H)**. The relationship remains specific to the post-inspiratory phase.

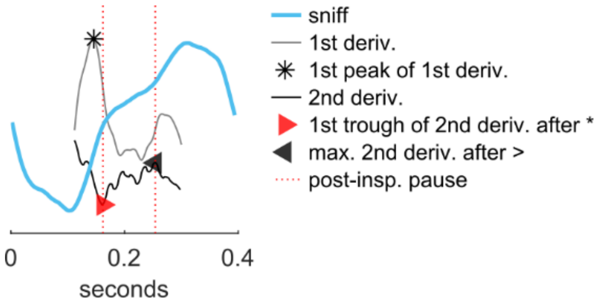


**Figure S6. Quantification of post-inspiratory pause.**
Method for detecting the post-inspiratory pause. The raw sniff trace (blue) is processed to calculate the first (gray) and second (black) derivatives. The pause onset (red triangle) is defined as the first trough of the 2nd derivative following the peak of the 1$^{st}$ derivative during inspiration (star), and the offset (black triangle) is defined as the subsequent peak of the 2nd derivative (see Methods).

### Table S1: Generalized Linear Mixed-Effects Model Results

**Model 1: Discrimination**

| Directionality: | Perf. Selection: | Observations (N): | DF: | | |
|---|---|---|---|---|---|
| Tu -> VTA | <0.8 | 70 | 64 | | |
| Formula: | | | | | |
| Channel Strength ~ 1 + Performance + Δ-Performance + Post-Insp. Pause + Unspecific Licking + Session # + (1 \| AnimalID) | | | | | |
| **Predictor** | **β (Estimate)** | **Std. Error** | **t-Stat** | **p-value** | **95% CI** |
| Intercept | -0.002 | 0.126 | -0.01 | 0.989 | [-0.254, 0.250] |
| Performance | 0.546 | 0.209 | 2.61 | 0.011 | [0.129, 0.963] |
| Δ-Performance | 0.007 | 0.113 | 0.06 | 0.953 | [-0.219, 0.233] |
| Post-Insp. Pause | 0.109 | 0.122 | 0.90 | 0.372 | [-0.134, 0.353] |
| Unspecific Licking | -0.451 | 0.212 | -2.13 | 0.037 | [-0.874, -0.028] |
| Session # | 0.143 | 0.126 | 1.14 | 0.259 | [-0.108, 0.395] |

**Model 2: Discrimination – directional specificity**

| Directionality: | Perf. Selection: | Observations (N): | DF: | | |
|---|---|---|---|---|---|
| VTA -> Tu | <0.8 | 70 | 64 | | |
| Formula: | | | | | |
| Channel Strength ~ 1 + Performance + Δ-Performance + Post-Insp. Pause + Unspecific Licking + Session # + (1 \| AnimalID) | | | | | |
| **Predictor** | **β (Estimate)** | **Std. Error** | **t-Stat** | **p-value** | **95% CI** |
| Intercept | 0.000 | 0.117 | 0.00 | 1.000 | [-0.234, 0.234] |
| Performance | 0.013 | 0.221 | 0.06 | 0.953 | [-0.429, 0.455] |
| Δ-Performance | 0.127 | 0.119 | 1.07 | 0.290 | [-0.111, 0.366] |
| Post-Insp. Pause | -0.030 | 0.128 | -0.23 | 0.815 | [-0.286, 0.225] |
| Unspecific Licking | 0.108 | 0.219 | 0.49 | 0.623 | [-0.330, 0.546] |
| Session # | -0.054 | 0.134 | -0.40 | 0.690 | [-0.321, 0.214] |

**Model 3: Discrimination – No-Go specificity**

| Directionality: | Perf. Selection: | Observations (N): | DF: | | |
|---|---|---|---|---|---|
| Tu -> VTA | <0.8 | 70 | 64 | | |
| Formula: | | | | | |
| Channel Strength ~ 1 + Performance (No-Go) + Δ-Performance + Post-Insp. Pause + Unspecific Licking + Session # + (1 \| AnimalID) | | | | | |
| **Predictor** | **β (Estimate)** | **Std. Error** | **t-Stat** | **p-value** | **95% CI** |
| Intercept | -0.000 | 0.115 | -0.00 | 1.000 | [-0.230, 0.230] |
| Performance (No-Go) | 0.033 | 0.198 | 0.17 | 0.867 | [-0.361, 0.428] |
| Δ-Performance | 0.010 | 0.118 | 0.09 | 0.932 | [-0.225, 0.246] |
| Post-Insp. Pause | 0.171 | 0.127 | 1.34 | 0.184 | [-0.083, 0.426] |
| Unspecific Licking | -0.011 | 0.208 | -0.05 | 0.957 | [-0.427, 0.405] |
| Session # | 0.204 | 0.131 | 1.56 | 0.124 | [-0.057, 0.466] |

**Supplemental Model 1: Shaping**

| Directionality: | Perf. Selection: | Observations (N): | DF: | | |
|---|---|---|---|---|---|
| Tu -> VTA | ~ | 50 | 44 | | |
| Formula: | | | | | |
| Channel Strength ~ 1 + Performance + Δ-Performance + Post-Insp. Pause + Unspecific Licking + Session # + (1 \| AnimalID) | | | | | |
| **Predictor** | **β (Estimate)** | **Std. Error** | **t-Stat** | **p-value** | **95% CI** |
| Intercept | -0.019 | 0.133 | -0.15 | 0.884 | [-0.287, 0.248] |
| Performance | 0.616 | 0.283 | 2.17 | 0.035 | [0.045, 1.186] |
| Δ-Performance | 0.158 | 0.151 | 1.04 | 0.302 | [-0.147, 0.462] |
| Post-Insp. Pause | -0.079 | 0.141 | -0.56 | 0.575 | [-0.363, 0.204] |
| Unspecific Licking | -0.646 | 0.291 | -2.22 | 0.031 | [-1.232, -0.061] |
| Session # | 0.106 | 0.149 | 0.71 | 0.479 | [-0.193, 0.406] |

**Supplemental Model 2: Discrimination – Go specificity**

| Directionality: | Perf. Selection: | Observations (N): | DF: | | |
|---|---|---|---|---|---|
| Tu -> VTA | <0.8 | 70 | 64 | | |
| Formula: | | | | | |
| Channel Strength ~ 1 + Performance (Go) + Δ-Performance + Post-Insp. Pause + Unspecific Licking + Session # + (1 \| AnimalID) | | | | | |
| **Predictor** | **β (Estimate)** | **Std. Error** | **t-Stat** | **p-value** | **95% CI** |
| Intercept | 0.001 | 0.128 | 0.01 | 0.991 | [-0.255, 0.258] |
| Performance (Go) | 0.564 | 0.250 | 2.26 | 0.028 | [0.065, 1.064] |
| Δ-Performance | 0.001 | 0.115 | 0.01 | 0.990 | [-0.227, 0.230] |
| Post-Insp. Pause | 0.151 | 0.121 | 1.25 | 0.216 | [-0.091, 0.394] |
| Unspecific Licking | -0.505 | 0.254 | -1.99 | 0.051 | [-1.012, 0.002] |
| Session # | 0.145 | 0.128 | 1.13 | 0.261 | [-0.110, 0.400] |
